# BML: a versatile web server for bipartite motif discovery

**DOI:** 10.1101/2021.05.28.446236

**Authors:** Mohammad Vahed, Majid Vahed, Lana X. Garmire

## Abstract

Motif discovery and characterization are important for gene regulation analysis. The lack of intuitive and integrative web servers impedes effective use of motifs. Here we describe **B**ipartite **M**otifs **L**earning (BML), a web server that provides a user-friendly portal for online discovery and analysis of sequence motifs, using high-throughput sequencing data as the input. BML utilizes both position weight matrix (PWM) and dinucleotide weight matrix (DWM), the latter of which enables the expression of the interdependencies of neighboring bases. With input parameters concerning the motifs are given, the BML achieves significantly higher accuracy than other available tools for motif finding. When no parameters are given by non-expert users, unlike other tools BML employs a learning method to identify motifs automatically and achieve accuracy comparable to the scenario where the parameters are set. The BML web server is freely available at http://motif.t-ridership.com/.

## INTRODUCTION

Transcription factors (TF) are essential regulatory patterns that control gene expression. Transcription factor binding sites (TFBS) are critical to a comprehension of gene expression regulations. Discovery of TFBS is one long-lasting issue in computational biology, with almost a hundred algorithms developed in the last 30 years _(Boeva V., 2016; Wasserman W. W., 2004; Sandve G. K., 2006). It is of importance not only to identify the binding motifs for a single block of TFBS but also motifs for two-block (bipartite) of TFBS separated by variable gaps. Many various types of bipartite motifs exist in both eukaryotes and prokaryotes _(Bi C. L., 2008). Position weight matrices (PWMs) are generally used to identify and represent TFBSs _(Stormo, 2000). They are based on the assumption that each nucleotide independently participates in the TFBSs of DNA sequence interaction. However, it has been long known that interactions between neighboring DNA bases affect TFBSs of DNA sequences interactions. For example, there are many interdependencies between neighboring positions of the binding sites of LexA and CRP in E. coli _(Salama, 2010). Methods that used dinucleotide weight matrix (DWM) outperformed those based on PWM _(Siddharthan, 2010).

Despite the algorithmic progress, most methods are difficult for non-experts to use. To overcome this problem, several web tools are available in the public domain, including MEME _(Bailey T. L., 2009), GLAM2 _Frith, 2008), Bipad _(Bi C. &., 2004; Lu R. M., 2017), Bioprospector _(Liu X. B., 2001), and AMD _(Shi, 2011). MEME and GLAM2 were developed for the prediction of one-block TFBS, while tools such as Bipad, Bioprospector, and AMD are designed for *ab initio* discovery of bipartite motifs for a set of DNA sequences. BioProspector is based on Gibbs sampling and BiPad is based on the entropy minimization algorithm. Both of them use PWM and enable bipartite motif prediction with variable gaps. On the other hand, AMD predicts bipartite motifs with constant gaps by comparing the target sequences with the background sequences. Most motif discovery web tools push certain levels of burden for motif parameterization to non-expert users, such as the motif length, direction, gap range, the number of occurrences, or the number of genes that include motifs. Some tools attempt to circumvent this problem by setting reasonable (yet not optimized) default parameters, however leading to sub-optimal performance.

To lessen the burden of motif parameterization for users but still deliver optimal performance, we propose using BML (short for **B**ipartite **M**otif **L**earning), a new parameter-free web server for motif-finding. BML is a novel ensemble learning method based on a hybrid of three-component algorithms (Gibbs sampling, Minimize Entropy, and Expectation-Maximization), each aimed at a specific category of motifs. BML can be used to find *de-novo* patterns, using input RNA or DNA sequences, beyond the prior knowledge of a database of known motifs. It is based on our previous Dipartite method _(Vahed, 2019), which discovers the bipartite motifs by considering the interdependencies of neighboring positions in the ChIP-seq data (Zhao, 2012). Specifically, we have improved BML in three aspects: (1) adding the graphical result such as sequence web logo and diagram; (2) increasing the accuracy of the results for the adjusted parameter method, and (3) adding a novel parameter-free method for discovering motifs.

## MATERIALS AND METHODS

### BML web server

We used ASP.NET, C#, Java, HTML, and CSS to implement BML. The absence of nuisance parameters enabled us to implement a clean and simple interface for input data, which can be either DNA or RNA sequences. Upon uploading or pasting sequence data, users can choose either with parameter mode, or parameter-free (PF) mode. User can select the method to find motif: PWM (Position Weight Matrix) or DWM (Dinucleotide Weight Matrix) that only require to set some parameters. In the parameter mode, users will need to provide some additional information, e.g., lengths of motifs and gaps, strand direction, option to allow degenerate sites, number of acceptable repeats, and the option of expect motif sites (One Occurrence Per Sequence, Any Number of Repetition, or Zero or One Occurrence Per Site). PF mode enables the practicing biologists to obtain results that comparable with those of other motif finders, without the hassle of guessing or experimenting the combination of parameters that gives the best performance. Methods supporting PF mode are described in the following sections.

Results from BML are displayed both in graphic and text formats. The best significant motifs found by BML are displayed graphically as consensus sequences on the main results page. Below it is a table containing summary statistics for each motif. Additional detailed information about the motifs is listed on the bottom part as texts. It shows the entropy scores in the iterations, consensus sequences, the position weight matrix (PWM, and DWM) constructed from all instances of the motifs and locations of the motif in each site (Figure 1).

**Fig 1.**
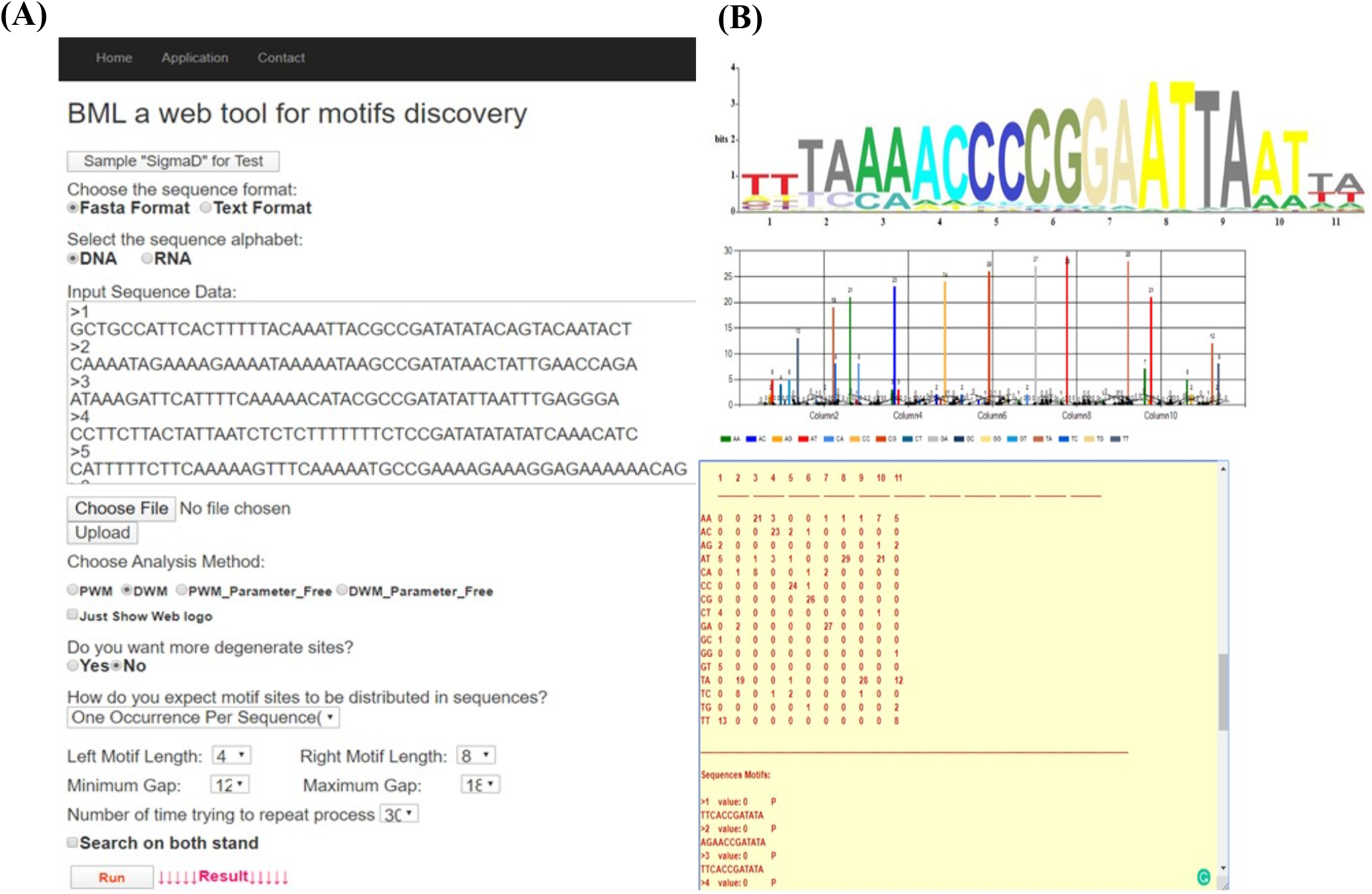
Snapshots of BML web-tool and outputs. (A) BML webtool takes the following inputs from users: sequencing data (in fasta or text format), DNA or RNA type, analysis method, and yes/no for degenerated sites. (B) After BML finishes the motif detection, the results of each prediction are shown graphically as Sequence Logo and text format including the position weight matrix diagram.

### Expectation-maximization expressions

The expectation maximization (EM) algorithm is a family of algorithms for learning probability models in problems that involve a hidden state _(Holmes, 2002). In our problem, the hidden state is where the motifs start in each training sequence. EM algorithm repeats the following two steps until convergence: ‘E’ step: Estimate the missing information using the current model parameters. ‘M’ step: Optimize the model parameters using the estimated missing information. The initial values for the motif model is done by randomly choosing the motif start point for each input sequence, then counts each nucleotide’s occurrence at different motif positions, and creates a consensus structure. The parameter-free mode implemented in BML assumes that each column in PWM or DWM can be identified as one nucleotide of the motif. In the context of motif discovery, this can be viewed as calculating the probability for the occurrences of the motif at certain position in the input dataset. The M-step then evaluates the estimation by maximizing the expected value of the log-likelihood function. Below, we describe EM algorithm mathematically.

Let’s denote the observed part of the data by X, and the missing information by Z. In our case, X represents the input sequences and Z the positions of the motifs. The aim is to find the model parameters θ maximizing the log-likelihood given the observed data:

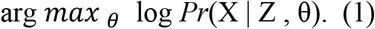

The EM algorithm is used to solve this optimization problem. The log-likelihood for the observed data might be difficult to obtain directly, thus we start by considering the conditional log-likelihood for the observed data given the missing information _(Lawrence, 1990; Gorodkin, 2011; Bailey T. L., The value of prior knowledge in discovering motifs with MEME., 1995; Bailey T. L., 2010; Bailey T. L., Fitting a mixture model by expectation maximization to discover motifs in bipolymers., 1994; Quang, 2014):

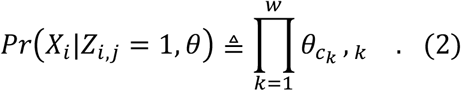

X_i_ is the ith sequence, and Z_i,j_ in the matrix Z represents the probability that the motif starts in position *j* in sequence *i*. The *w* is the length of the motif, *c* is the length of each sequence, and *k* is position of the motif, 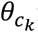 represents the probability of character *c* in column k. We calculate the probability of a training sequence given a hypothesized starting position:

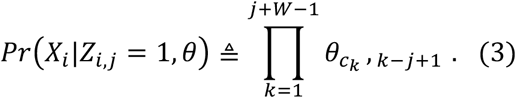

To calculate the probability of a motif, BML computes the probability of motif sequence range, as well as the probability of these background sequences before and after the motif range. It is the outcome of probabilities over the *w* positions in the motif and the remaining background positions (Supplementary Figure S1). this time using background probabilities for all positions within sequence i:

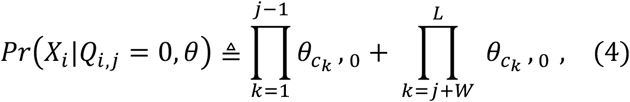

Q functions as the {X, Z} background log-likelihood. The novel index variable Q_i,j_ is 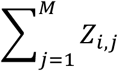 and *j* is motif start position.

In the E-step, the anticipated counts of all nucleotides at each position are calculated based on the parameters’ current guess. The E-step needs the assessment of the probability of the hidden data p(Z|X, θ), that is, 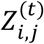 for each position. Then, Bayes’ theorem is applied to specify 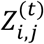 in parts of (6):

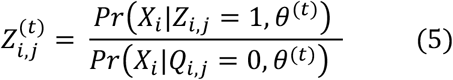

For all *i* ∈ {1, …, *N*} and *j* ∈ {1, …, *M*}. (*N* is # of sequences, *M* is # of Nucleotides in each sequence) In the M-step, all of parameters are updated based on the values calculated in the E-step. The M-step estimating *θ*, recall *θ*_*c,k*_ represents the probability of character c in position:

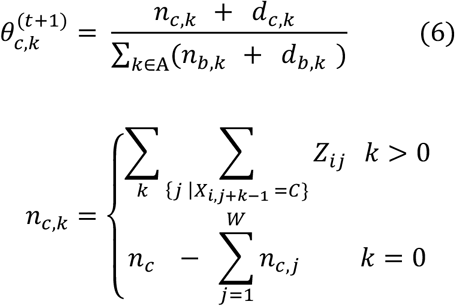

For *j* ∈ {1, …, W}, *A* ∈ {A, C, G, T} for PWM and *A* ∈ {AA, AC, AG, AT, CA, …, TT} for DWM, *d* is pseudo-counts ({*d*_A_, *d*_C_, *d*_G_, *d*_T_} *or* {*d*_AA_, *d*_AC_, *d*_AG_, … *d*_TT_}), and *n*_*c*_ is total # of *c*’*s* in data set and *b* is background.

BML iteratively does the E and M steps until the change in formula (Euclidean distance) falls below a threshold (default: 10^−8^). EM algorithm makes maximum likelihood estimation to maximize an objective function. Since the E and M steps are repeated, the EM algorithm converges to a maximum.

### Objective function

The objective function minimizes Shannon’s entropy for PWM and DWM of the concatenated TFs of the left and right motifs, in Equation 7 below:

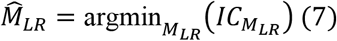

where *M*_*LR*_ is the concatenated motif, and 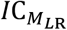 is the entropy for the motif *M*_*LR*_. 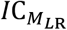 is given by:

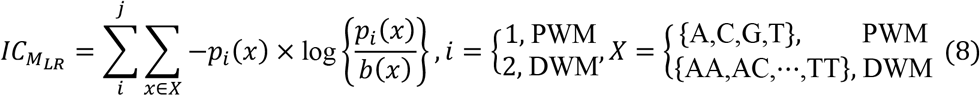

Where *x* is a mononucleotides in PWM, or a dinucleotide for DWM, *p*_*i*_ (*x*) and *b*(*x*) are the compositions of *x* in the motif sites and the background sites (no motif sites), respectively. *j* is the sum of the lengths of the left and right motifs. *p*_*i*_(*x*) and *b*(*x*) are given by:

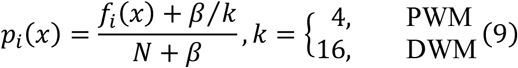

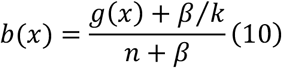

where *f*_*i*_(*x*) is the frequency of *x* at position *i*, i.e., the mononucleotide at the position *i* for PWM (or PWM-L), or the dinucleotide at the positions *i* − 1, and *i* for DWM (or DWM-L). *β* is the pseudo-count (*β*=1). *k* is the number of the patterns, i.e., *k* = 4 for PWM (PWM-L) or *k* = 16 for DWM (DWM-L). *N* is number of sequences in the input data. *g*(*x*) is the frequency of *x* in the background sites. *n* is the total number of the background mononucleotides for PWM (PWM-L) or background dinucleotides for DWM (DWM-L), which do not harbor the motif sites.

### Filtering motifs

To reduce the False Positive scores and optimize motif results, we add the cut-off value on the standardized motif scores. The normalization is done as the following:

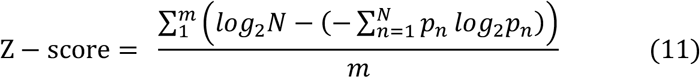

m is number of sites in input sequences data, *p*_*n*_ is the detected frequency of symbol *n* at a special sequence position, and N is the number of different symbols for the read sequence type. As a result, the maximum sequence conservation per site is log_2_ 4 = 2 bits for Dinucleotide (DWM) and log_2_ 16 = 4 bits for Mononucleotide (PWM). Only motifs with scores larger than the minimum cut-off scores (equation 12) are shown as output of BML.

### Real Data

#### Cyclic AMP receptor protein (CRP) data

CRP is a prokaryotic TF that is significant in regulating genes involved in energy metabolism. It binds to 22-bp consensus motif sites in Escherichia coli. The sequences were recovered from Regulon Database as “TFBSs” (Release: 9.4 Date: 05-08-2017) _(Gama-Castro, 2016). Among 374 sequences of CRP binding sites, we used 323 unique sequences for performance comparison whose lengths are 36 - 42 bp. The motif lengths are: 16 bp (11 sites), 17 bp (1 site), 20 bp (1 site), 22 bp (308 sites), and 23 bp (two sites).

#### Promoter motifs in human

We selected the motifs among 1460 motifs in human _(Xie, 2005), where the left and right motifs (or two-block motifs) are more than 3-nt and the gap length is more than the left/right motif length, due to the limitation of Bioprospector and Bipad in detecting bipartite motifs. As a result, we obtained 40 motifs. We retrieved the promoter sequences around each binding site (500 bp upstream to 500 bp downstream) as the datasets.

GLAM2 has an input size limit of 60000 bp and did not work for 11 of the 40 datasets, namely, CCCNNNNNNAAGWT-4, GGCNNNNNKCCAR-13, GTTNMNNNNNAAC-18, GTTNNNNNKNAAC-19, MCAATNNNNNGCG-23, MYAATNNNNNNNGGC-25, TGGNNNNNNKCCAR-32, TTTNNNNNAACW-35, WGTTNNNNNAAA-38, YKACANNNNNCAGA-39, YTGGMNNNNNGCC-44, YTGGMNNNNNNCCA-45.

### Sigma factor data

We used nine datasets of bipartite motifs with variable gap lengths from the sigma factor dataset in *B. subtilis* from DBTBS _(Makita, 2004). Minimum and maximum gap lengths, and left motif and right motif lengths are determined by DBTBS, with the abbreviation Motif_Left_ <(Min_Gap_,Max_Gap_]>Motif_Rigth_ :

-SigA (344 sequences ranging between 38 bp to 93 bp, 6<[11,23]>6).

-SigB (64 sequences ranging between 39 bp to 64 bp, 6<[12,18]>6).

-SigD (30 sequences ranging between 44 bp to 57 bp, 4<[12,18]>8).

-SigE (70 sequences ranging between 41 bp to 58 bp, 7<[12,18]>8).

-SigF (25 sequences ranging between 41 bp to 71 bp, 5<[13,19]>10).

-SigG (55 sequences ranging between 40 bp to 76 bp, 5<[15,20]>7).

-SigH (25 sequences ranging between 41 bp to 60 bp, 7<[9,18]>5).

-SigK (53 sequences ranging between 38 bp to 85 bp, 4<[9,17]>9).

-SigW (34 sequences between from 38 bp to 53 bp, 10<[13,17]>6).

### Other programs used for comparison

Five popular tools, namely MEME (ver. 5.1.1), GLAM2 (ver. 5.1.1), BioProspector (release 2), BiPad (ver. 2), and AMD, were compared with BML. For the CRP dataset, MEME was executed with the options “-mod oops”, “-dna”, “-minw 22”, “-maxw 22”, and “-w 22”. GLAM2 was executed with the options “-z 100” “-a 22” “-b 22” “-w 22” “-r 1” “-n 2000” “-D 0.1” “-E 2.0” “-I 0.02” “-J 1.0”. BioProspector was executed with the options “Width (22, 22)”, “Gap (0, 0)”, “-n 50”, and “-n 3”. BiPad was executed with the options “-l 22”, “-a 0”, “-r 0”, “-i”, “-b 0”, and “-y 1000”. AMD was executed with the options “-MI” and “-T 1”. For human promoter and Sigma factor datasets, we used the same above settings for each tool, but different motif size and gap ranges. AMD did not work on SigE and SigF, we used option “-T 2”. For the background sequences in AMD, we used the 200 bp upstream areas of 4,314 genes in *E. coli* K-12 (NC_000913.3) for CRP TF binding sequence data, the promoter sequences of human genes (hg17: upstream1000.fa.gz) for human promoter TF binding sequence dataset, and the 200 bp upstream areas of 4,448 genes in *B. subtilis* 168 (NC_000964.3) for Sigma factors.

### Evaluation of prediction accuracy

To evaluate the performance of each tool, we use the nucleotide-level correlation coefficient (nCC) with the same datasets and parameters _(Tompa, 2005). The nCC is calculated as:

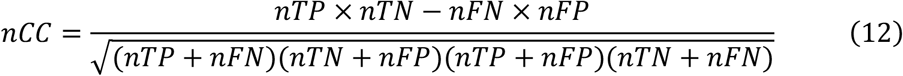

nTP is the number of nucleotides in sequences that correctly detected motifs, nFP is the number of background nucleotides incorrectly detected motifs, nTN is the number of background nucleotides correctly identify as the background, 4. nFN is the number of nucleotides with motifs but incorrectly be called as the background. The graphic illustrations of nTP, nFP, nTN, and nFN are provided in Supplementary Figure S2.

## Results

### Summary of BML Web Server

We used ASP.NET, C#, Java, HTML and CSS to implement BML. In designing the web interface for BML, we aimed for the optimal user experience with as little technical nuisance as possible, maximized amount of output information, and minimalized run-time for each task. BML takes sequence data as the input and uses two methods to find motifs: PWM (Position Weight Matrix) and DWM (Dinucleotide Weight Matrix). It also has two modes of motif discovery: with parameters or parameters-free (PF) mode. In the parameter mode, the users can choose the default values or decide a small set of input parameters, such as length of motifs and gaps, sequence type as DNA or RNA, forward or reverse direction, option to allow degenerate motif sites, the number of iterations to run the algorithm, and expected motif site distribution in the sequences (Figure 1A). In the PF mode, users do not need to set any parameters, instead the BML-PF methods (PWM-PF and DWM-PF) will predict the motifs based on sequence input data alone.

Results from BML are displayed both in the graphic and text format (Figure 1B). BML shows the best motif logos graphically on the main results page, with a table containing summary statistics for each motif. It also shows more detailed information about the motif in the text format underneath, such as the entropy scores over iterations, the position weight matrix (PWM, and DWM) constructed from all instances of the motifs, and the starting/ending site of each motif. A typical run of BML takes between 5 seconds ∼ 10 minutes. Runtime is dependent mostly on the size of the used input data and whether users provide parameter values in BML. BML-PF method is slower, due to its search space over a wide number of parameters.

### Benchmarking BML (with parameters) against other web tools

Using experimentally determined datasets as the testing cases, BML identified motifs with a significantly higher accuracy level than several other popular tools motif finders (with their default parameters). Below we describe the comparison results on CRP TF binding sequence data, human promoter sequence data, and sigma factor data in bacteria *B. subtilis* 168 sequentially. We use nucleotide-level correlation coefficient (nCC) as the metric for accuracy (see Methods).

#### Performance on CRP sequence dataset

We evaluated the performance of BML (BML_PWM and BML_DWM) with the five popular motif discovery tools (MEME, GLAM2, BioProspector, BiPad, and AMD) by using the TF binding sites of CRP (Figure 2A-B). We chose 323 unique sequences out of 374 sequences of CRP binding sites as the test datasets. For testing the one-block motif, namely, the 22-bp motif, BML performs the best among all methods. BML_DWM is slightly better than BML_PWM, as expected, with a nCC of 0.944 (Figure 2A). Next we compared the accuracy on detecting the bipartite motif on these sequences (Figure 2B). Similar to the annotation in Dipartite [14], we annotate a biparitite motif as: L<d>R where L and R are the lengths of left and right motifs respectively, and d is gap range. Among all three types of bipartite motifs 8<[6]>8, 6<[8]>8, 6<[10]>6, BML_PWM and BML_DWM are superior to BiPad and BioProspector. To evaluate the robustness of the methods, we randomly sampled 100 datasets with 100 sequences from these CRP binding sites. Again we obtained consistent conclusion: BML_DWM and BML_PWM slightly outperform other tested tools (Supplementary Figure S3). Taking them together, BML shows better performance at identifying the bipartite and one-block motifs.

**Fig 2.**
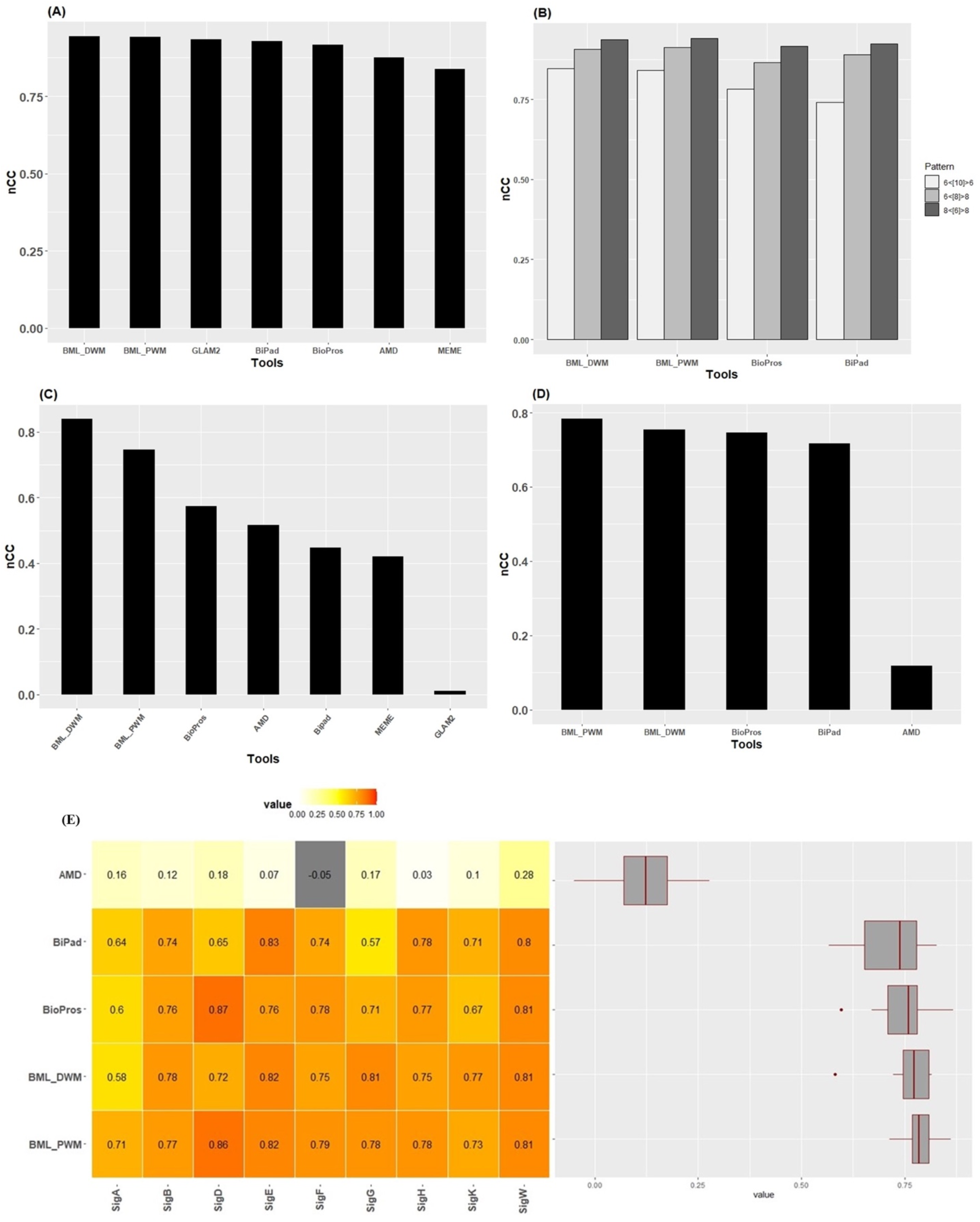
The motif detection comparison on CRP sequences, Human Promoter, and sigma factor dataset in *B. subtilis*. (A) Average *nCC* results for searching the one-block motif, i.e., the 22 bp motif, in descending order on BML webserver (BML_PWM and BML_DWM modes), GLAM2, BiPad, BioProspector, AMD, and MEME. (B) Average *nCC* results for searching the bipartite motifs, i.e., 6<[10]>6, 6<[8]>8, and 8<[6]>8, on BML_PWM, BML_DWM, BioProspector, BiPad, and PWM. (C) The combined *nCC* values calculated using a total of 3,054 sequences from 40 bipartite human motifs datasets. Note: GLAM2 can only support 29 datasets due to the limit of max input sequence length of 60000 bp. (D) Average nCC results of all sigma (σ) factor datasets. Datasets σA, σB, σD, σE, σF, σG, σH, σK, and σW consist of 344, 64, 30, 70, 25, 55, 25, 53, and 34 sequences, respectively. (E) Heat map and Boxplots of the *nCC* values by different motif methods on all 9 Sigma datasets.

#### Performance on human promoter sequence datasets

We evaluated the performance of bipartite motif detection by different methods on the 40 human promoter motif datasets that met our selection criteria (see Methods). Due to the input size limit of GLAM2, it only accepted 29 out 40 datasets. BML_DWM significantly outperforms other tested tools on the human promoter datasets, with the highest nCC=0.84 and is followed by BML_PWM with nCC=0.74 (Figure 2C). This confirms that DWM improves the bipartite motif detection upon PWM, as the dataset does consist of dinucleotides. Other tools Bioperspector, AMD, Bipad, MEME, and GLAM2 have sequentially descending performance after BML methods, with nCCs of 0.57, 0.44, 0.42, and 0.01 respectively. Closer examinations show that AMD, Bipad, and MEME also have larger interquartile ranges, suggesting the lack of stability in their predictions (Supplementary Figure S4).

#### Performance on sigma factor datasets

We next evaluated the performance of BML with other methods over bipartite motifs with variable gaps between two block motifs. For this, we used 9 bipartite motifs in sigma TF in *B. subtilis*, from DBTBS as the testing datasets. The 9 sigma factor datasets are, SigA (344 sequences), SigB (64 sequences), SigD (30 sequences), SigE (70 sequences), SigF (25 sequences), SigG (55 sequences), SigH (25 sequences), SigK (53 sequences), and SigW (34 sequences). Overall BML methods (BML_PWM and BML_DWM) perform the best, with the average nCC of 0.78 and 0.75 for BML_PWM and BML_DWM respectively (Figure 2D and 2E). They also show the smallest variations among datasets (Figure 2E). AMD has significantly worse nCC values (average nCC=0.11) than all other methods, confirming that it is not desirable at handling variable gap lengths in bipartite motifs.

### Using BML to predict bipartite motifs with the parameter-free (PF) mode

In some cases, the users do not know what best parameters should be used as the input. To handle these situations, they may use BML-PF mode (Figure 3) to find the bipartite motifs with variable gaps based on input sequences only. The BML-PF algorithm is inspired by unsupervised approach to estimate the parameters of the probability distributions to best fit the motifs of a given dataset. It is implemented by an iterative approach of EM algorithm that cycles between two steps (Equations 5 and 6, Methods). The EM algorithm is applied quite widely in machine learning and is frequently used in unsupervised learning problems, such as density estimation and clustering without input parameters _(Toivonen, 2018; Figueiredo, 2002; Bailey T. L., Unsupervised learning of multiple motifs in biopolymers using expectation maximization., 1995; Yang, 2011). In our application, the first step (E-step) attempts to estimate the latent variables. The second step (M-step) attempts to optimize the parameters of the model to discover the best motifs on sequences data (Figure 3B).

**Fig 3.**
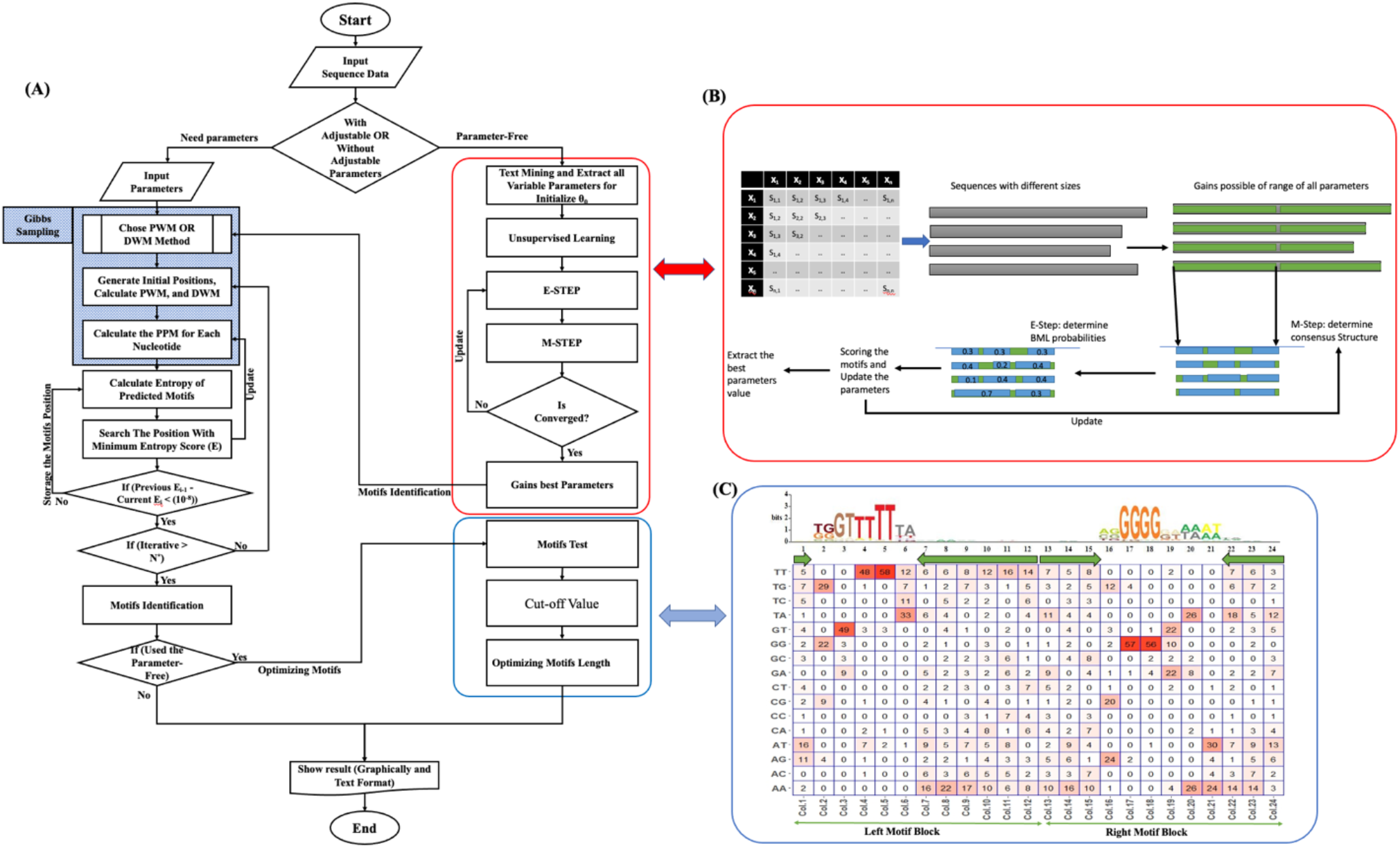
Illustration of BML algorithm. (A) The overall workflow of BML. BML uses two methods, with parameter and parameter free modes. It proposes bipartite motifs based on PWM or DWM iteratively. Each iteration starts from randomly generated positions until differences of the entropy are minimized. (B) Strategy of EM algorithm for BML_PF to extract the best parameter values. Initialized with the maximumly acceptable motif length, in each iteration EM algorithm updates Position Probability Matrix (PPM) and improves the positions of the motifs by M-Step and E-Step, then updates the motif score table. (C) Normalization and filtering of motif sequences in BML_PF. In this step dynamic programming is used to update the motif scores. The low-value motif sequences are filtered separated from the beginning and the end of motifs body.

Initialized with the maximumly acceptable motif length, in each iteration EM algorithm updates Position Probability Matrix (PPM) and improves the positions of the motifs (Figure 3B). The sequences data along with the best fit parameters extracted in the EM step are sent back to the Gibbs sampling for motif predictions (Figure 3A). We have observed that the EM algorithm sometimes chooses some nucleotides with low scores as a motif. To reduce the False Positive scores and optimize results, we add the cut-off value on the standardized motif scores. Only motifs with scores larger than the minimum cut-off scores (Equation 11, see Methods) are shown as output (Figure 3C). As a default, BML assumes 0.5 as the minimum Z-score shown in the sequence logo.

### Comparing PF and parameter-based modes in BML

We first demonstrate BML results in the parameter-free (PF) mode as compared to the set-parameter mode (Figure 4), on the same CRP TF binding sequence, Human promoter sequence, and Sigma factor datasets as described earlier. We first evaluated the effect of input data size on the performance of BML-PF using the sequences of SigA in *B. subtilis* (Figure 4A-B). By randomly sampling the sequences of SigA, we generated 100 datasets, where each dataset contains 10, 20, 50, 100, 150, 200, and 300 sequences respectively. With increasing the size of the datasets, BML exhibits better performance on both the average values and variances of nCC scores (Figure 4A-B). BML_PWM-PF variances for the datasets with 200 and 300 sequences are relatively close to those of BML_PWM (Figure 4A). Interestingly, BML_DWM-PF has even better performance than BML_DWM on this simulated dataset based on SigA (Figure 4B). On the other hand, the averaged results of BML parameter-free method (BML_PWM-PF and BML_DWM-PF) are both similar but slightly less compared to BML with adjusted parameters (BML_PWM and BML_DWM), on both CRP TF binding (Figure 4C) and on all 9 sigma factor datasets (Figure 4D). Out of 9 sigma factors, SigW has the highest average nCCs (Figure 1E) in both PWM (0.808) and DWM (0.811) approaches, indicating the presence of base interdependencies in the motif of SigW. We thus compared the PF and with parameter modes to the real motif in SigW next (Figure 4E). BML PWM_PF and DWM_PF have nCCs of 0.752 and 0.808 respectively, very similar to the scenarios with parameters where PWM and DWM approaches have nCCs of 0.808 and 0.811 respectively. This shows the accuracy of the BML-PF method. Confirming by the logos, all 4 approaches largely recover the bipartite motif as determined by DBTBS, which includes 10 bp (TGAAACTTT) on the left and 6 bp (CGTATA) on the right.

**Fig 4.**
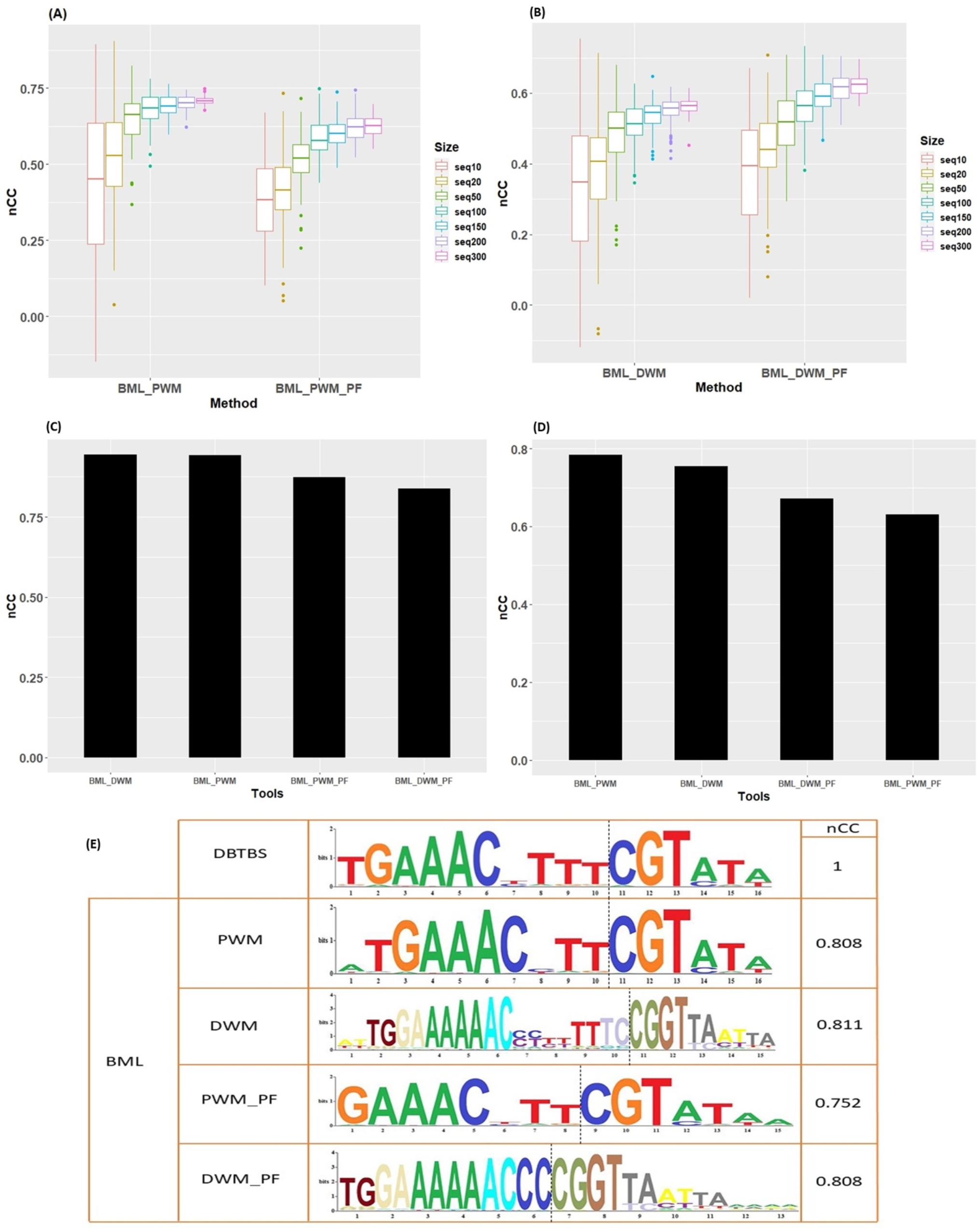
The performance of BML in PF mode. (A-B) The robust assessment on randomly generated subsets from SingA dataset using PF modes. 100 datasets were generated by sub-sampling of the SigA dataset, yielding 10, 20, 50, 100, 150, 200, 300 sequences in each of the 100 datasets. (A) nCCs of BML_PWM and BML_PWM_PF; (B) nCCs of BML_DWM and BML_DWM_PF. (C) Average nCC results for searching the one-block motif of CRP TF binding sequences. (D) Average nCC results for searching the one-block motif of all 9 sigma factor datasets. (E) Sequence logos of Motifs discovered on SigW dataset, by BML using PWM and DWM with parameterization, or under PF mode using PWM_PF and DWM_PF.

## CONCLUSION

Here we present BML web server, a very user-friendly motif exploration method that enables both the expert and non-expert biologists to identify one-block motifs and bipartite motifs. We evaluate the performance of BML on various datasets compared with other freely available tools, namely, MEME, GLAM2, BioProspector, AMD, and BiPad for motif discovery. We show that BML performs significantly better than these alternatives. When naïve users are not sure about the values of the input parameters, they can use BML as a parameter-free web server and still retrieve consistent motifs. BML is available for use at http://motif.t-ridership.com/.

## AUTHORS’ CONTRIBUTION

MV envisioned the project and conducted the implementation and analysis, MV participated in designing and coordinating the study. LXG supervised the study and provided funding support. MV and LXG wrote the manuscript. All authors have read and approved the manuscript submission.

## ACKNOWLEDGEMENT

This research was supported by grants K01ES025434 awarded by NIEHS through funds provided by the trans-NIH Big Data to Knowledge (BD2K) initiative (www.bd2k.nih.gov), P20 COBRE GM103457 awarded by NIH/NIGMS, R01 LM012373 awarded by NLM, R01 HD084633 awarded by NICHD to L.X. Garmire.

## Supplementary

**Supplementary Figure S1.**
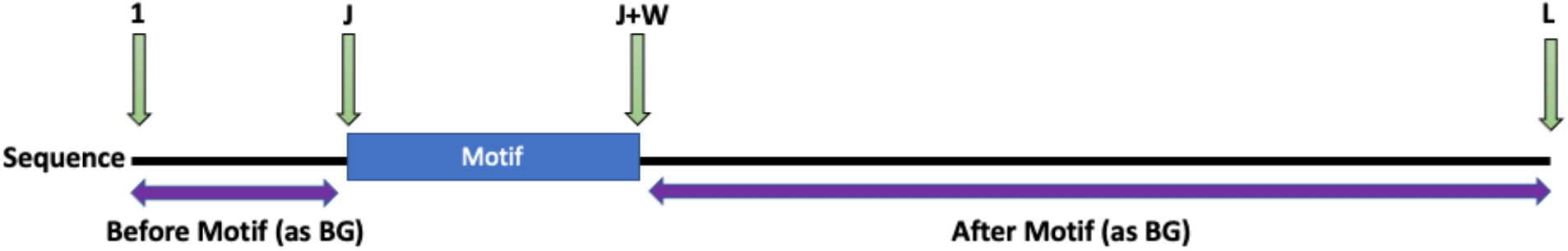
Illustration of the position of a motif and the background sequences.

**Supplementary Figure S2.**
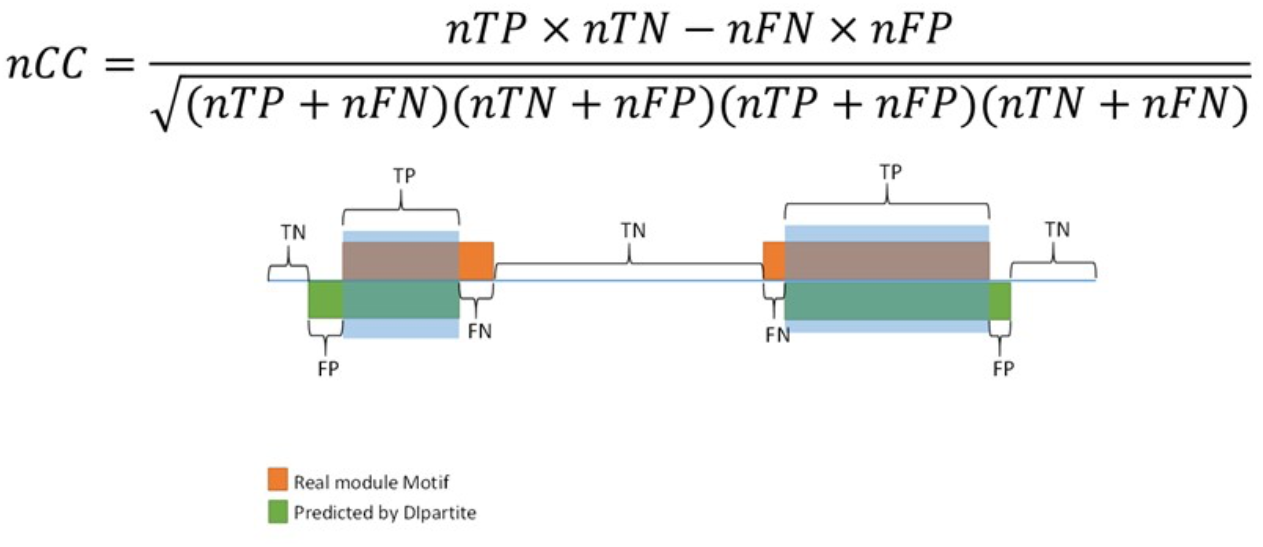
The formular to calculate nucleotide-level correlation coefficient (nCC), the metric of accuracy in Motif discovery. nCC is given by:

TP: is the number of nucleotide positions in both known sites and predicted sites.

TN: is the number of nucleotide positions in known sites but not in predicted sites.

FP: is the number of nucleotide positions not in known sites but in predicted sites.

FN: is the number of nucleotide positions in neither known sites nor predicted sites.

**Supplementary Figure S3.**
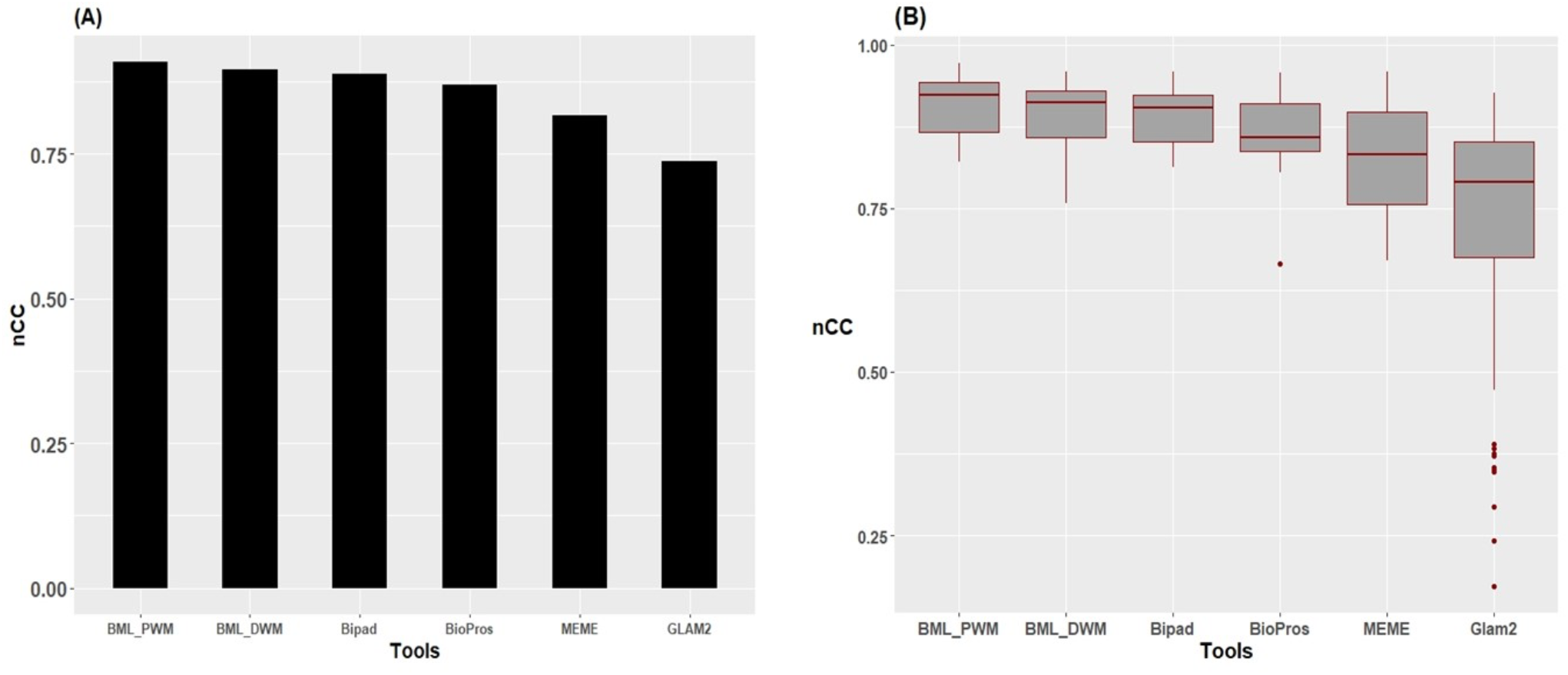
The performance comparison on 100 CRP TF binding site datasets from searching the one-block motif (22 bp). 100 datasets consisting of 100 sequences were generated by randomly sampling the CRP datasets. (A) Average *nCC* results in descending order on BML webserver (BML_PWM and BML_DWM modes), BiPad, BioProspector (BioPros), MEME and GLAM2. (B) Boxplots of the same results in (A).

## Notes

### Competing Interest Statement

The authors have declared no competing interest.

